# The genome sizes of ostracod crustaceans correlate with body size and phylogeny

**DOI:** 10.1101/114660

**Authors:** Nicholas W. Jeffery, Emily A. Ellis, Todd H. Oakley, T. Ryan Gregory

**Affiliations:** Department of Integrative Biology, University of Guelph, Guelph, Ontario, Canada. N1G 2W1; University of California Santa Barbara, Santa Barbara, California, USA. 93106

**Author notes:** Present address: Fisheries and Oceans Canada, Bedford Institute of Oceanography, Dartmouth, Nova Scotia (Nicholas W. Jeffery).

**Keywords:** Ostracod, genome size, C-value, body size, phylogeny

## Abstract

Within animals a positive correlation between genome size and body size has been detected in several taxa but not in others, such that it remains unknown how pervasive this pattern may be. Here we provide another example of a positive relationship, in a group of crustaceans whose genome sizes have not previously been investigated. We analyze genome size estimates for 46 species across Class Ostracoda, including 29 new estimates made using Feulgen image analysis densitometry and flow cytometry. Genome sizes in this group range ~80-fold, a level of variability that is otherwise not seen in crustaceans with the exception of some malacostracan orders. We find a strong positive correlation between genome size and body size across all species, including after phylogenetic correction. We additionally detect evidence of XX/XO sex determination in all three species of myodocopids where male and female genome sizes were estimated. On average, genome sizes are larger but less variable in myodocopids than in podocopids, and marine ostracods have larger genomes than freshwater species, but this appears to be explained by phylogenetic inertia. The relationship between phylogeny, genome size, body size, and habitat is complex in this system, and will benefit from additional data collection across various habitats and ostracod taxa.

## Introduction

Genome sizes (haploid nuclear DNA contents) have been estimated for more than 5,500 species of animals, revealing a greater than 7,000-fold range (Gregory 2017). Some important patterns have emerged from comparative analyses of genome size, including well-established links between genome size, nucleus size, cell size, and cell division rate (Gregory, 2002; Gregory, 2005). However, the implications of these relationships at organismal and ecological levels are more complex. Body size (Gregory, Hebert and Kolasa, 2000; Jeffery, Yampolsky and Gregory, 2016; Wyngaard, *et al.,* 2005), metabolic rate (Gregory, 2002; Vinogradov, 1995), developmental rate (Gregory, 2002; Gregory and Johnston, 2008), life history (Rees, *et al.,* 2007; Reeves, *et al.,* 1998), and geographic distribution (Bonnivard, *et al.,* 2009) have been found to correlate with genome size in some animal groups, but these are far from universal, and depend on the biology of the animals in question. Sorting out the broader patterns of genome size diversity and its biological significance therefore requires studies of a wide variety of taxa. However, many groups -- especially non-vertebrates -- remain to be studied from this perspective.

Ostracod crustaceans represent a group of interest in this regard, due to their ecological diversity and over 150-fold range in body size. Class Ostracoda contains an estimated 20,000 species divided into four orders, the two largest of which are Myodocopida and Podocopida. About 10% of extant ostracod species are freshwater, while the remainder are marine (Horne, Cohen and Martens, 2002). Myodocopids are strictly marine, but podocopids can be found in marine, estuarine, and freshwater habitats, with three podocopid lineages having invaded nonmarine habitats independently at different times since the Devonian (Martens, *et al.,* 2008). The ostracod clade has a rich fossil record (Foote and Sepkoski, 1999) dating to at least the early Ordovician (Williams, *et al.,* 2008), but molecular evidence suggests an origin at least 500 MYA, while divergence of Myodocopida and Podocopida occurred approximately 480 MYA (Oakley, *et al.,* 2013; Tinn and Oakley, 2008).

Prior to this study, there were only two published ostracod genome size estimates, with an additional 15 unpublished values included in the Animal Genome Size Database and (P.D.N. Hebert unpublished). Genome size estimates are sparse for crustaceans in general, and obtaining a greater number of estimates for ostracods will provide data for potential genome sequencing projects and provide a better understanding of how a basic organismal property such as genome size can influence the overall life history of these crustaceans.

Here, we report new estimates for 29 species of ostracods from two orders - Myodocopida and Podocopida - and analyze them in combination with publicly available estimates. We also constructed a genus-level phylogeny for the taxa with genome size estimates using 18S rRNA sequences to examine genome size diversity across the ostracod phylogeny. This allowed us to test possible relationships between genome size and body size within a phylogenetic context, and to examine patterns of genome size distribution across taxa and habitats.

## Methods

### Specimen collection and biological/ecological trait data

We collected sediment samples from various locations in Canada, the United States, and Australia (Table 1, Supplementary Table S1). To collect marine and freshwater species, we used a 150 μ m plankton net to horizontally collect the top half-inch layer of sediment from the benthos. The resulting sediment from marine samples was further sorted using a 500 μ m sieve and a 100 μ m sieve, and sorted by eye using a dissection microscope. We collected some species at night – as many are known to be nocturnally active in search of mates (Speiser, *et al.,* 2013) – by using a light trap and vertical plankton tows. In many cases, we were able to identify taxa to the species level using published generic and specific dichotomous keys. We measured total body size (length of carapace) using an ocular micrometer.

**Table 1.**
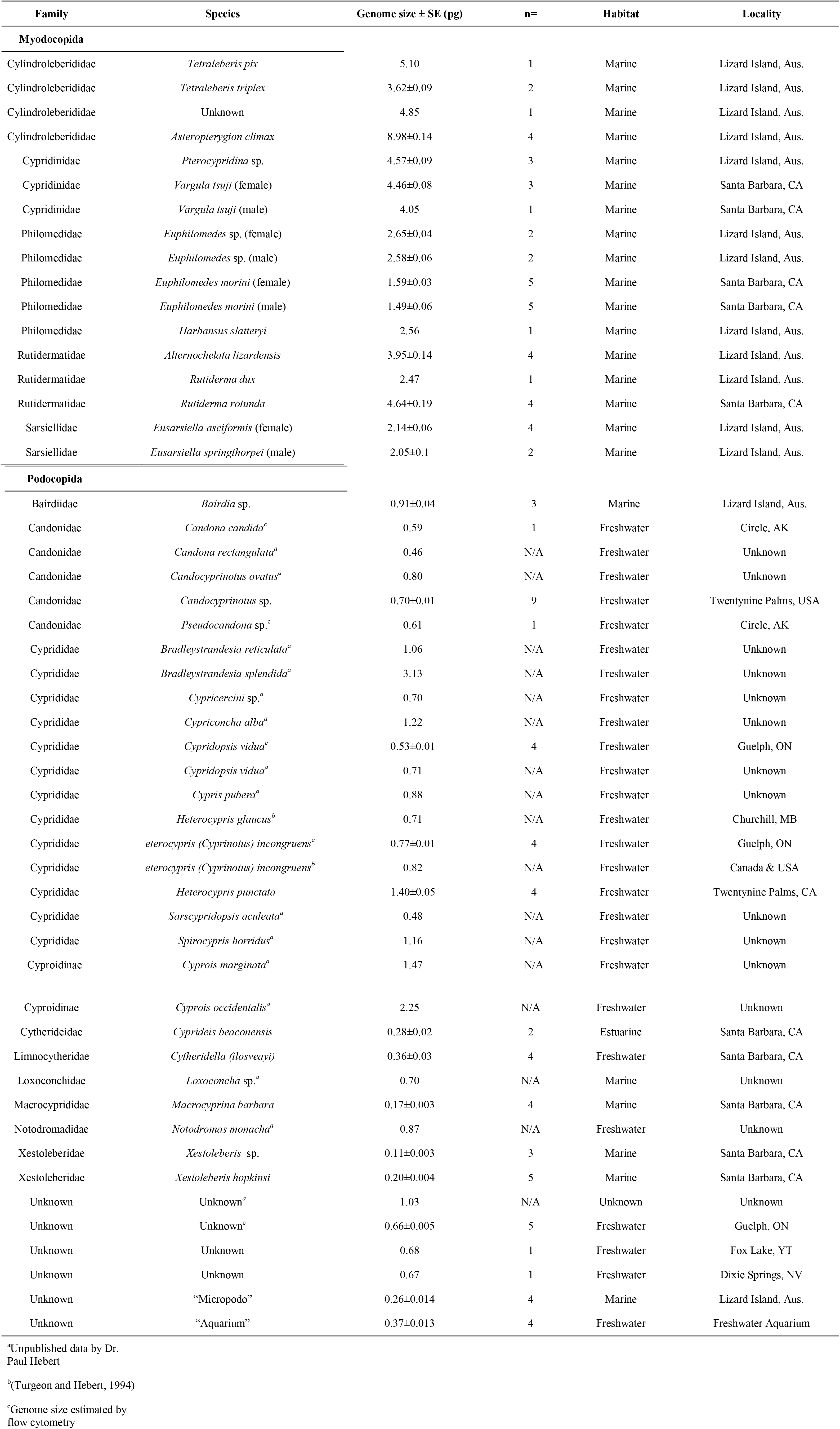
Genome size estimates in picograms for 46 species of ostracods from the orders Myodocopida and Podocopida collected from Canada, Australia and the USA. When known, the sex is also listed after the species name. The data includes published values for two species as well as unpublished data by P. Hebert.

### Genome size estimation

Ostracods were dissected from their valves in 40%(v/v) acetic acid on microscope slides using insect pins, flattened with a coverslip which was held onto the slide with a clothespin, and then frozen on dry ice. The coverslip was then removed with a razor blade and the slides were immersed in 100% ethanol and dried at room temperature.

All prepared slides were fixed in 85:10:5 methanol:formalin:acetic acid overnight. The slides underwent hydrolysis in 5N HCl at 20°C for 120min. The slides were stained in prepared Schiff reagent for 120min, followed by 3 rinses in bisulfite solution and repeated rinses in deionized water (Hardie et al. 2002). Each set of slides was co-stained with slides of domestic chicken blood (1C=1.25pg) and rainbow trout blood (1C=2.60pg) for use as internal standards. Individual nucleus densities were measured using a Leica DM LS compound microscope with a mounted Optronics DEI-750 CE CCD camera and Bioquant Life Science software.

We estimated the genome size of some species by flow cytometry (Table 1). Specimens were crushed in 500μl cold LB01 buffer (Dolezel, Binarova and Lucretti, 1989) and co-stained with chicken blood or crushed *Daphnia pulex* (1C=0.23pg) after adding 12μl propidium iodide (24μg/ml) and 2μl RNase (4μg/ml) to each sample. These were stained in the dark for 1 hour and analyzed on an FC500 flow cytometer (Beckman-Coulter). All coefficients of variation (CVs) were <8% and a minimum of 1000 nuclei per ostracod were analyzed.

We included 17 estimates from the Animal Genome Size Database (http://www.genomesize.com) (Hebert, unpublished). These estimates used epithelial tissue dissected from ostracods, but the standard used for estimation is unknown.

### Statistical analyses and phylogenetic correction

We first applied a general linear model of log-transformed genome size, log-transformed body size, and habitat type as a categorical variable (0=freshwater, 1=marine) to test for significant relationships between variables without phylogenetic correction. To correct for phylogenetic non-independence of our data, we used software implemented in the publically available Galaxy computing platform Osiris (Oakley, *et al.,* 2014), to estimate a genus-level phylogeny from publicly available 18S ribosomal RNA sequences (Supplementary Table S2). These sequences were aligned using MUSCLE v3.8 (Edgar, 2004). We performed maximum-likelihood and bootstrap statistical analyses in Garli v2.0 (Gutell and Jansen, 2006) using a GTR+G model with 8 gamma categories. For tips in the phylogeny for which we had multiple species per representative genus in the phylogeny, we took the average genome size. The resulting phylogeny had 16 genera of ostracods and relationships were consistent with previously published phylogenies (Oakley, *et al.,* 2013; Tinn and Oakley, 2008) (Figure 1). We used the package *phytools* (Revell, 2012) in R v3.2.3 to map genome size onto our phylogeny and conduct a test for phylogenetic signal using Pagel’s lambda. We then conducted a phylogenetic generalized least squares model of genome size, body size, and habitat, using lambda as the correlation structure, and implemented in the R packages *ape* (Paradis, Claude and Strimmer, 2004) and *nlme* (Pinheiro, *et al.,* 2014).

**Figure 1.**
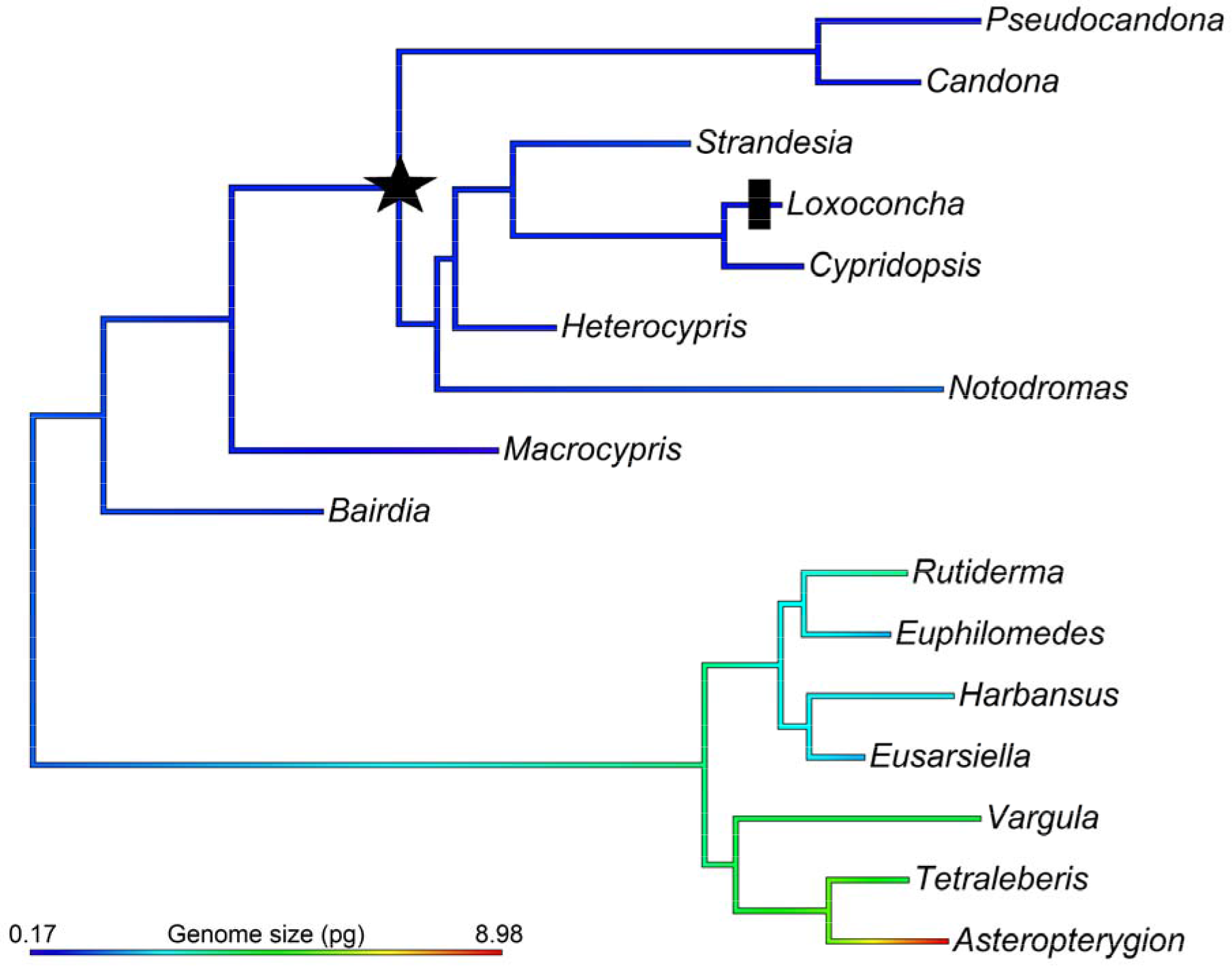
Genus-level 18S phylogeny showing podocopid and myodocopid clades with genome size mapped onto the branches. The transition to freshwater is marked with a star, and the reversal to marine is marked with a black bar.

## Results

Genome size estimates for all analyzed ostracods range approximately 80-fold, from 0.11±0.003pg (mean±s.e.) in the podocopid *Xestoleberis* sp. to 8.98±0.14pg in the myodocopid *Asteropterygion climax* (Figure 1). Freshwater ostracods, found entirely in the order Podocopida, had an average genome size of 0.86±0.09pg, while marine ostracods from both Podocopida and Myodocopida had an average genome size of 2.88±0.46pg. The single estuarine species in this study, *C. beaconensis,* had a small genome size of 0.28±0.02pg. Larger genomes are found within Myodocopida on average (3.81±0.51 pg) versus the Podocopida (0.82±0.11pg); however, the Podocopida exhibit a higher coefficient of variation in genome size (73.6%) relative to Myodocopida (49.8%). For three species within Myodocopida, we estimated genome size for both males and females, and found that males have lower DNA content by an average of 6.3% (Table 1).

A linear regression of non-phylogenetically corrected log-transformed genome size versus body size showed a significant positive relationship across all of the species for which genome sizes were estimated in this study (r^2^=0.47, p<0.0005, n=41) (Figure 2), and a general linear model of genome size, body size, and habitat revealed that both body size and habitat were significantly associated with genome size (t=6.351 and p<0.0001, t=3.076 and p=0.004 respectively). When splitting ostracods by order, both myodocopids and podocopids showed a significant relationship between transformed genome size and body size (r^2^ =0.53, p=0.002 and r^2^=0.31, p=0.001 respectively) (Figure 2). To examine this relationship within a phylogenetic context, it was necessary to conduct the analysis at the genus level (n=15 genera). After controlling for phylogeny, body size remained significantly associated with genome size (t=3.63, p=0.004) but habitat did not (t=-0.45, p=0.66). Overall, genome size showed moderate but significant phylogenetic signal for the genera examined here (λ=0.60, p=0.027).

**Figure 2.**
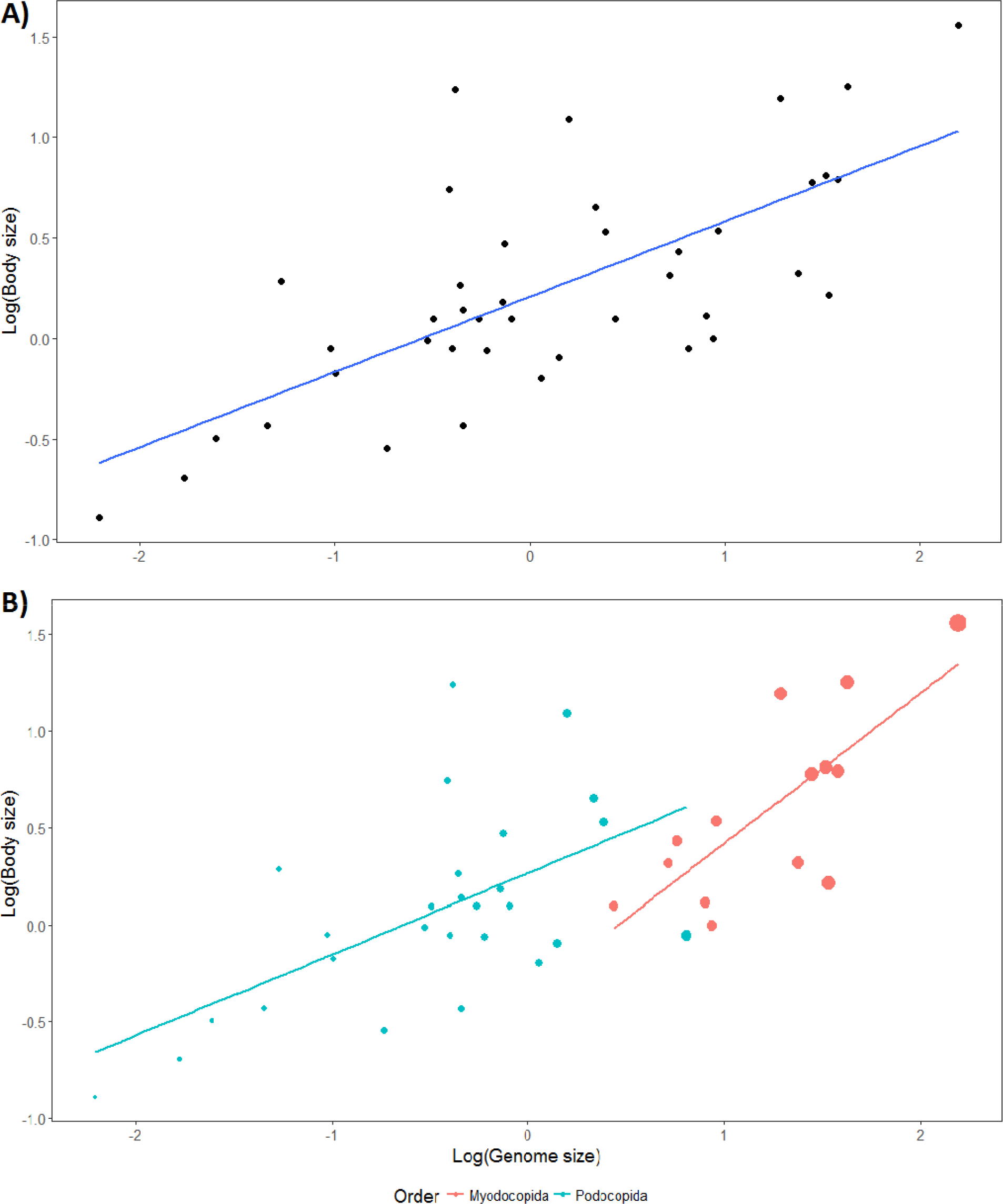
Log-transformed body size versus log-transformed genome size in (A) 41 species of ostracods showing a significant positive relationship (r^2^=0.47, p<0.0005) and (B) splitting the relationship by order reveals a significant relationship within both Myodocopida and Podocopida (r^2^=0.53, p=0.002 and r^2^=0.31, p=0.001 respectively). Dot size is proportional to genome size in panel B) to show the larger genomes in Myodocopida overall.

## Discussion

### Genome size in ostracods

We revealed an 80-fold range in genome size within ostracods, a large range which has not been observed in other crustaceans with the exception of Isopoda and Amphipoda (Jeffery, 2015). Our genus-level phylogeny is largely consistent with previously published phylogenies (e.g. Yamaguchi and Endo, 2003) and we find moderate but significant phylogenetic signal for genome size across the phylogeny. This suggests that genome size evolution deviates from a pure Brownian motion model, which would be expected if λ=1.0, and rather that genome size correlates modestly among closely related species and differs between more deeply divergent clades. This could be consistent with evidence for potential polyploidy or quantum leaps in genome size across this phylogeny, such as the case between the sister-genera *Tetraleberis* and *Asteropterygion,* where the genome of *Asteropterygion* is nearly double that of *Tetraleberis.* Discontinuous patterns of genome size have been reported in other crustaceans, including copepods (e.g. Gregory, *et al.,* 2000).

### Genome size, body size, habitat, and sex

Larger genomes are observed in the order Myodocopida, which have larger body sizes than the Podocopida. Moreover, the positive relationship between genome size and body size remained even after correcting for phylogeny. This positive relationship between body size and genome size has only been observed in few other crustaceans, including copepods (Gregory, *et al.,* 2000; Wyngaard and Rasch, 2000), amphipods, and some branchiopods (Hessen and Persson, 2009). This correlation suggests that body size is a good predictor of genome size in ostracods, though this is by no means an exhaustive examination of this trend. It remains to be seen how general the body size correlation is within and among crustacean groups (and other animals) more broadly. It is likely that this relationship will be most evident when cell numbers contribute less to body size diversity than individual cell sizes.

Habitat initially was revealed to be significantly correlated with genome size, as larger genomes appear to be found in marine habitats. However after correcting for phylogeny, habitat remained no longer significant. This is because Myodocopida are entirely marine and have larger genome sizes than the predominantly freshwater Podocopida, but within Podocopida the marine species we examined had smaller genomes than freshwater species. Denser taxon sampling could certainly change these conclusions.

We revealed that in the three myodocopid species for which we had individuals of each sex, males had smaller genome sizes than their female counterparts. This is consistent with an XX♀/XO♂)' sex determination system in which males lack one copy of a sex chromosome that is present in two copies among females (Moguilevsky, 1990). The extent to which this sex determination system occurs across ostracods is unknown, but we detected evidence in Families Cypridinidae and Philomedidae, and previous research has found evidence in each of these families (Moguilevsky, 1985; Moguilevsky, 1990; Rivera and Oakley, 2009). This suggests that the common ancestor of all myodocopids had XX/XO sex determination (Figure 1).

### Alternative hypotheses of genome size evolution

We observe here that genome size is highly variable among ostracods and that genome size correlates with body size, but are unable to determine if these patterns are the result of ecological pressures leading to rapid genome size change in extant species. Podocopids possess much smaller genomes than myodocopids on average, suggesting that they are subject to additional constraints on genome size, perhaps relating to body size or other ecological pressures that impact cell size and/or cell division rate. Indeed, the myodocopids tend to be physically larger in size than podocopids. It is also noteworthy that the Myodocopida are strictly marine, whereas the Podocopida inhabit a much broader range of habitats (leaf litter, freshwater, brackish, marine) (Martens, *et al.,* 2008). This could help to explain why, despite small absolute genome sizes, the Podocopida exhibit higher relative variability in genome size due to differential selective pressures in different environments, as reflected by coefficient of variation. While on average we find larger genomes in marine species, a simple marine versus freshwater distinction is not likely to account for genome size differences across ostracods, however, given that within the Podocopida, marine species tend to have smaller genomes than freshwater species. Within Ostracoda, genome size may thus be driven by differential selective pressures for changes in body size within lineages and across habitat types.

### Concluding remarks

The present study has highlighted some interesting patterns in genome size diversity among the previously overlooked Ostracoda. We provided new genome size estimates for 29 species of ostracods relative to only 17 estimates contained within the Animal Genome Size Database. Even with a comparatively small sampling of overall ostracod diversity, a more than 80-fold range in genome size was observed. This, combined with the relationships between genome size and body size, and potential differences in selective pressures from different habitats indicate that this is an excellent group to target for further study.

## Authors’ Contributions

NWJ and TRG conceived of the study. NWJ, EAE, and THO collected and identified specimens. NWJ and EAE prepared the samples and EAE constructed the phylogeny. NWJ and EAE analyzed the samples and results. NWJ, EAE, THO, and TRG wrote the paper. All authors edited and approved of the paper.

## Acknowledgements

We are thankful to Dr. Paul Hebert for allowing us to include his unpublished ostracod genome size data. We are also grateful to Alexandra Wen and Abigail Chua for their assistance in collecting and preparing slides for genome size analysis. We thank the Australian Museum (loan number MI2013059) for loaning us prepared slides and Paul Valentich-Scott for assistance in accessioning slides (accession numbers x-x) at the Santa Barbara Museum of Natural History.

## Funding

NWJ and TRG are supported by the National Sciences and Engineering Research Council of Canada (Canada Graduate Scholarship and Discovery Grant, respectively). This research was funded through National Science Foundation Grant #DEB-1146337 to THO, Research Mentorship Program funding to A. Wen and EAE, and Australian Museum Grant #SB14014 to Celia K.C. Churchill.

## Data Availability

Sampling locations and species identifications are available in Table 1. All 18S sequence Genbank accession numbers are provided in Supplementary Table S1. All genome size estimates will be publicly available in the Animal Genome Size Database prior to final publication.

